# Native MOWChIP-seq: Genome-wide profiles of key protein bindings reveal functional differences among various brain regions

**DOI:** 10.1101/2021.07.11.451963

**Authors:** Zhengzhi Liu, Lynette B. Naler, Yan Zhu, Chengyu Deng, Qiang Zhang, Bohan Zhu, Zirui Zhou, Mimosa Sarma, Alexander Murray, Hehuang Xie, Chang Lu

## Abstract

Genome-wide profiling of interactions between genome and various functional proteins is critical for understanding regulatory processes involved in development and diseases. Conventional assays require a large number of cells and high-quality data on tissue samples are scarce. Here we optimized a low-input chromatin immunoprecipitation followed by sequencing (ChIP-seq) technology for profiling RNA polymerase II (Pol II), transcription factor (TF), and enzyme binding at the genome scale. The new approach, termed native MOWChIP-seq, produces high-quality binding profiles using 1000-50,000 cells. We used the approach to examine the binding of Pol II and two TFs (EGR1 and MEF2C) in cerebellum and prefrontal cortex of mouse brain and found that their binding profiles are highly reflective of the functional differences between the two brain regions. Our analysis reveals the potential for linking genome-wide TF or Pol II profiles with neuroanatomical origins of brain cells.

## Introduction

Protein-DNA interactions are widely and critically involved in the regulations of gene transcription and expression^1^. RNA polymerase II (Pol II) transcribes protein-coding genes into mRNA following several steps: formation of preinitiation complex, promoter-proximal pausing, elongation and termination. Pol II binding occurs throughout the genome, with enrichment at regions being either actively expressed or readied for imminent transcription upon environmental cues^2,3^. This latter phenomenon of Pol II pausing is the rate limiting step on more than 70% of metazoan genes and has been shown to play key biological roles.^3–6^ Similarly, transcription factors (TFs) bind to DNA or cofactors and participate in gene regulations in a sequence-specific manner.^1,7^ TFs control key aspects of cellular biology including cell differentiation, development patterning, and immune response. ^8,9^ Finally, enzymes such as histone acetyl transferases (HATs) and histone deacetylases (HDACs) closely interact with the genome to effect epigenetic modifications (histone acetylations) that are critically involved in gene silencing and activation^10^. For example, HDAC2 binding and activity are involved in memory formation and synaptic plasticity in brain^11^. Genome-wide binding profiles of these proteins are highly indicative of genes being activated or readied in a process. Difference in the binding profile among various tissue samples reveals underlying genome-wide molecular dynamics and variations involved in development or disease.

Chromatin immunoprecipitation coupled with sequencing (ChIP-seq) is a simple and direct approach to profile *in vivo* genome-wide binding of proteins. Although ChIP-seq has been generating reliable and high-quality data on histone modifications (i.e. the interaction between a modification histone and the genome)^12–14^, ChIP-seq results on other types of protein-DNA interactions tend to be much more challenging. Compared to the robust histone-DNA interaction, the interaction between DNA and other protein molecules such as Pol II, TFs, and enzymes may be harder to preserve even after treatment such as formaldehyde crosslinking. Thus conventional ChIP-seq requires a large quantity of starting material (>10^7^ cells per assay) for probing genomic binding of proteins that are not histones and the results tend to have lower reproducibility compared to histone ChIP-seq.^15^ These limitations makes it impractical to examine animal and human tissue samples available only in small quantities. In recent years, significant efforts have been made in developing low-input ChIP-seq methods^12,14,16–22^. However, the vast majority of these methods were only been demonstrated on examining histone modifications.

As an alternative to ChIP, IP-free technologies such as CUT&RUN^23,24^ and ChIL-seq^25^ were developed to profile factor binding to the genome. CUT&RUN maps Pol II and TF binding by cutting and releasing DNA fragments that interact with antibody-targeted Pol II and TFs into supernatant, requiring as few as 1,000 cells ^23,24^. The latest variation of CUT&RUN (CUT&Tag)^26^ have been used to profile single cells. However, CUT&RUN requires immobilization of cells to bead surface via concanavalin A and glycolipid interaction and CUT&RUN data appear to exhibit lower correlation with the gold-standard ChIP-seq data (e.g. ENCODE data) compared to low-input ChIP-seq technologies ^23,24^. CUT&RUN datasets may also be contaminated by DNA sequences that are from unknown sources associated with the process (e.g. sequences containing (TA)_n_)^27^.

Here we establish micrococcal nuclease (MNase)-digestion-based native MOWChIP-seq as a general tool for low-input profiling of protein binding to genome. MNase digestion was previously applied in native ChIP to probe mostly histone modifications^28^ and also non-histone proteins including pol II and TFs^29–33^. However, these studies were all conducted using a large number of cells (10^7^−10^8^). Our approach, referred to as native MOWChIP-seq or nMOWChIP-seq, combines MNase digestion of native chromatin with a microfluidics-based low-input ChIP-seq technology (MOWChIP-seq^12,14^) that was previously applied to examine histone modifications only. We show that MNase digestion is effective for preserving the links between protein and genome and facilitating low-input profiling. We generated high-quality ChIP-seq data using as few as 1,000 cells for studying Pol II, 5,000 cells for TF EGR1, and 50,000 cells for HDAC2. We applied this method to study genome-wide binding of Pol II, EGR1 and MEF2C in two functional regions of the mouse brain: prefrontal cortex (PFC) and cerebellum. Extensive variations in RNA Pol II and TF binding were identified between these two brain regions and these profiles reveal the involvement of these regulatory molecules in the functional difference.

## Results

### Profiling genome-wide binding of RNA Pol II, TFs, and enzymes

RNA Pol II, TF or enzyme binding has been conventionally studied after crosslinking using reagent such as formaldehyde that firmly immobilizes the protein to the genomic DNA ^34,35^. However, our initial results showed that even using our low-input MOWChIP-seq technology which has proven high efficiency for collecting DNA-protein complexes^12,14^, low-quality results were yielded when 100,000 cells were crosslinked and sonicated to create the chromatin fragments before Pol II ChIP (Supplementary Fig. S1). 54,000 and 17,000 peaks were yielded in the two technical replicates, compared to 115,000 and 141,000 peaks generated by nMOWChIP-seq using 10,000 cells per assay. Crosslinking and subsequent sonication may damage RNA Pol II’s ability to form antigen-antibody complex and seriously limits the ChIP’s efficiency for collecting targeted fragments.

We then applied nMOWChIP-seq to profile protein binding to the genome. MNase digestion (or native ChIP) had primarily been applied to studies associated with histone modifications where the strong interaction between genomic DNA and histone guarantees the pulldown of DNA without crosslinking ^13,23,36^. In our process (Fig. 1A), tissues were first mechanically homogenized to extract nuclei and cultured cells were directly used. Nuclei or cells were lysed and digested with MNase to yield chromatin fragments with size range appropriate for ChIP (150-600 bp). That was followed by the MOWChIP process^12,14^. Briefly, in a microfluidic chamber with a partially closed sieve valve, antibody-coated magnetic IP beads were packed into a dense bed. All chromatin fragments were forced through the bead bed, drastically increasing the adsorption efficiency of targeted fragments. Oscillatory washing was then applied to remove non-specific binding, and the beads with bound chromatins were transferred out of the chamber for DNA elution, library preparation and sequencing.

**Fig 1.**
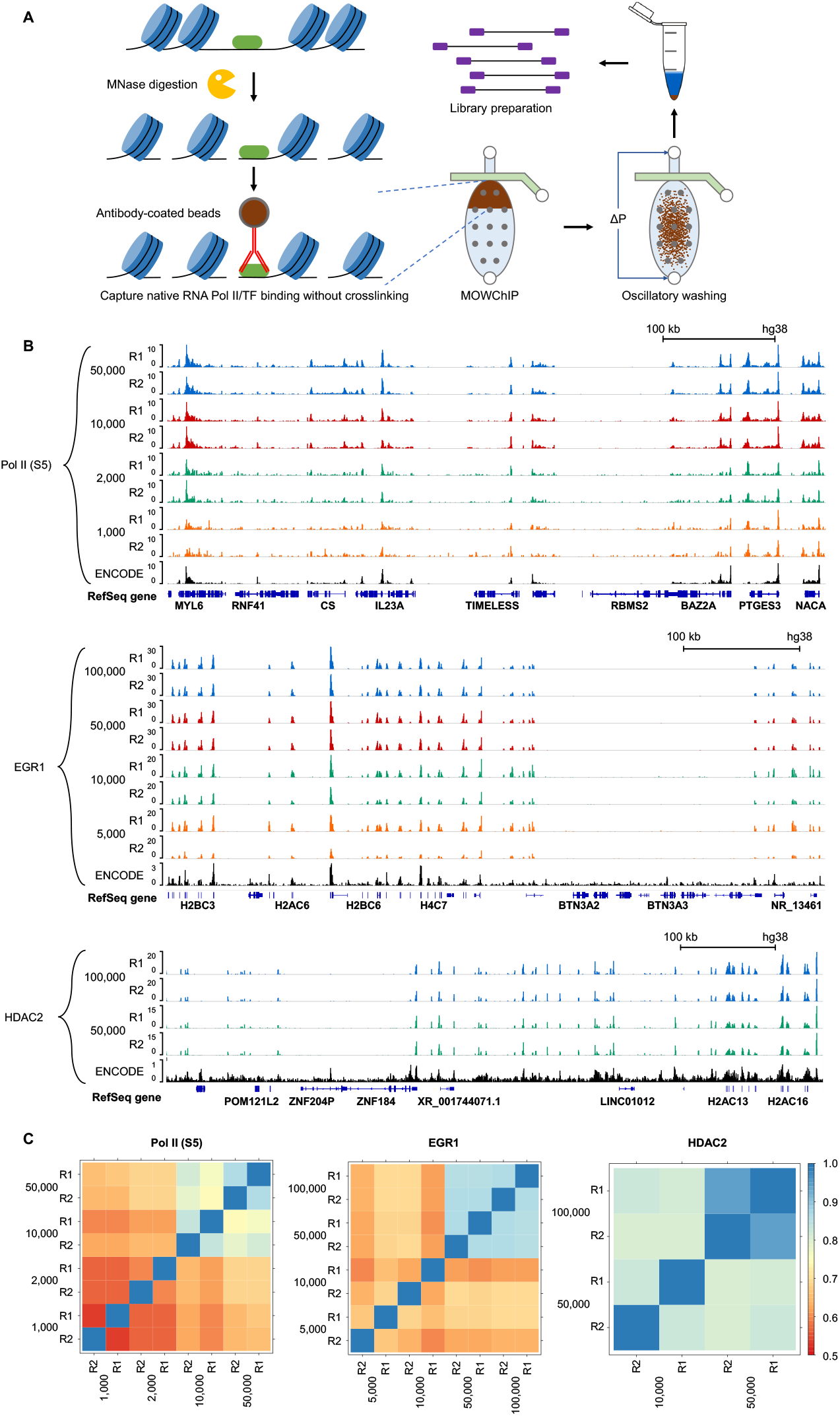
Overview of the low-input nMOWChIP process and data on RNA polymerase II, EGR1 and HDAC2 binding in GM12878 cells. (A) Steps to generate nMOWChIP-seq libraries. (B) Normalized RNA Pol II (S5) signals generated using 1,000 to 50,000 cells, EGR1 signals generated using 5,000 to 100,000 cells, and HDAC2 signals generated using 50,000 and 100,000 cells. ENCODE data (Pol II: GSM803485; EGR1: GSM803434; HDAC2: GSE105521) are included for comparison. (C) Pearson’s correlations of affinity score calculated from consensus MACS2 peaks and uniquely mapped reads using DiffBind, among samples of various cell numbers for RNA Pol II, EGR1 and HDAC2.

Using cell line GM12878, we show that our method can generate high quality ChIP-seq data with input as little as 1,000 cells per assay for Pol II (phospho S5) (Fig. 1B). Pearson correlations between technical replicates were 0.87, 0.84, 0.60, and 0.54 for 50,000-, 10,000-, 2,000-, and 1,000-cell samples on RNA Pol II, respectively (Figure 1C). We benchmarked our data against ENCODE data obtained using conventional ChIP-seq method and 20 million cells per assay. We examined the number of peaks called and the fraction of reads in peaks (FRiP) for data quality (Supplementary Table S1). The nMOWChIP-seq Pol II data produced averagely 137,000, 128,000, 65,000 and 27,000 peaks with 50,000, 10,000, 2,000 and 1,000 cells per assay, respectively, compared to 110,000 peaks from the ENCODE data obtained using 20 million cells. FRiP measures the amount of background in the data and nMOWChIP-seq Pol II data showed a decrease from 49.8% to 9.4% with cells per assays decreasing from 50,000 to 1,000, far exceeding the 1% threshold recommended by ENCODE^37^.

We also determined that the binding of Pol II was stable enough to be preserved at −80 °C, a unique property not observed in histone modifications or TFs (Supplementary Figure S2). From our testing, MNase-digested chromatin can be frozen at −80 °C or on dry ice for 2 d without substantial degradation in data quality for Pol II.

We profiled a transcription factor, early growth response protein 1 (EGR1), using as few as 5,000 GM12878 cells (Fig. 1B and 1C, Supplementary Table S1). Pearson correlation coefficients between the technical replicates were 0.87, 0.87, 0.63, and 0.67 for 100,000-, 50,000-, 10,000-, and 5,000-cell samples, respectively (Fig. 1C). EGR1 data generally show lower peak numbers and FRiP than Pol II data, as shown by both nMOWChIP-seq and ENCODE data. FRiP ranged from 8.4% to 3.2% in our nMOWChIP-seq EGR1 data. We also profiled another TF MEF2C (Supplementary Table S1). A large percentage of the EGR1 (85%) and MEF2C (75%) peaks overlap with Pol II peaks (Supplementary Figure S3).

Finally, we applied nMOWChIP-seq to examine the binding of histone deacetylase HDAC2 in GM12878 cells (Fig. 1B and 1C, Supplementary Table S1). At least 50,000 cells were required to generate good quality data on HDAC2 binding. An average of 1,394 (FRiP = 1.2%) and 8527 peaks (FRiP = 3.1%) were generated using 50,000 and 100,000 cells per assay, compared to 1820 peaks of ENCODE data obtained using 10 million cells (FRiP = 0.3%). We also examined HDAC2 binding in mouse brain cells (Supplementary Fig. S4). 100,000 mixed nuclei and 800,000 FACS-sorted NeuN+ neuronal nuclei were used in each assay and an average of 3193 (FRiP ~13.1%) and 2112 (FRiP ~10.4%) peaks were produced (Supplementary Table S1).

### Comparison of Pol II and Pol II-S5 binding profiles

We used two antibodies to differentiate the binding of phosphor-5 activated subset of Pol II (Pol II-S5, ab5131) and all Pol II regardless of its phosphorylation status (Pol II-total, ab817, clone 8WG16). Our data show the distinction (Fig. 2). The two profiles on GM12878 cells are similar in a large fraction of the genome (Fig. 2A). However, some genes (ACTG1 and MDM2 as examples) with Pol II binding over the entire gene body show that Pol II-total profile captures the initial binding of unphosphorylated Pol II at TSS, which is absent in Pol II-S5 profile in comparison (Fig. 2B). During the course of binding to a gene, Pol II is unphosphorylated during the pre-initiation stage, but undergoes phosphorylation once a short (20-60 bp) mRNA begins to transcribe.^2^ Being able to differentiate the distinct forms of Pol II is important when the exact Pol II binding status on specific genes is of interest (e.g. whether pre-initiation or activated Pol II dominates pausing at a TSS).

**Fig 2.**
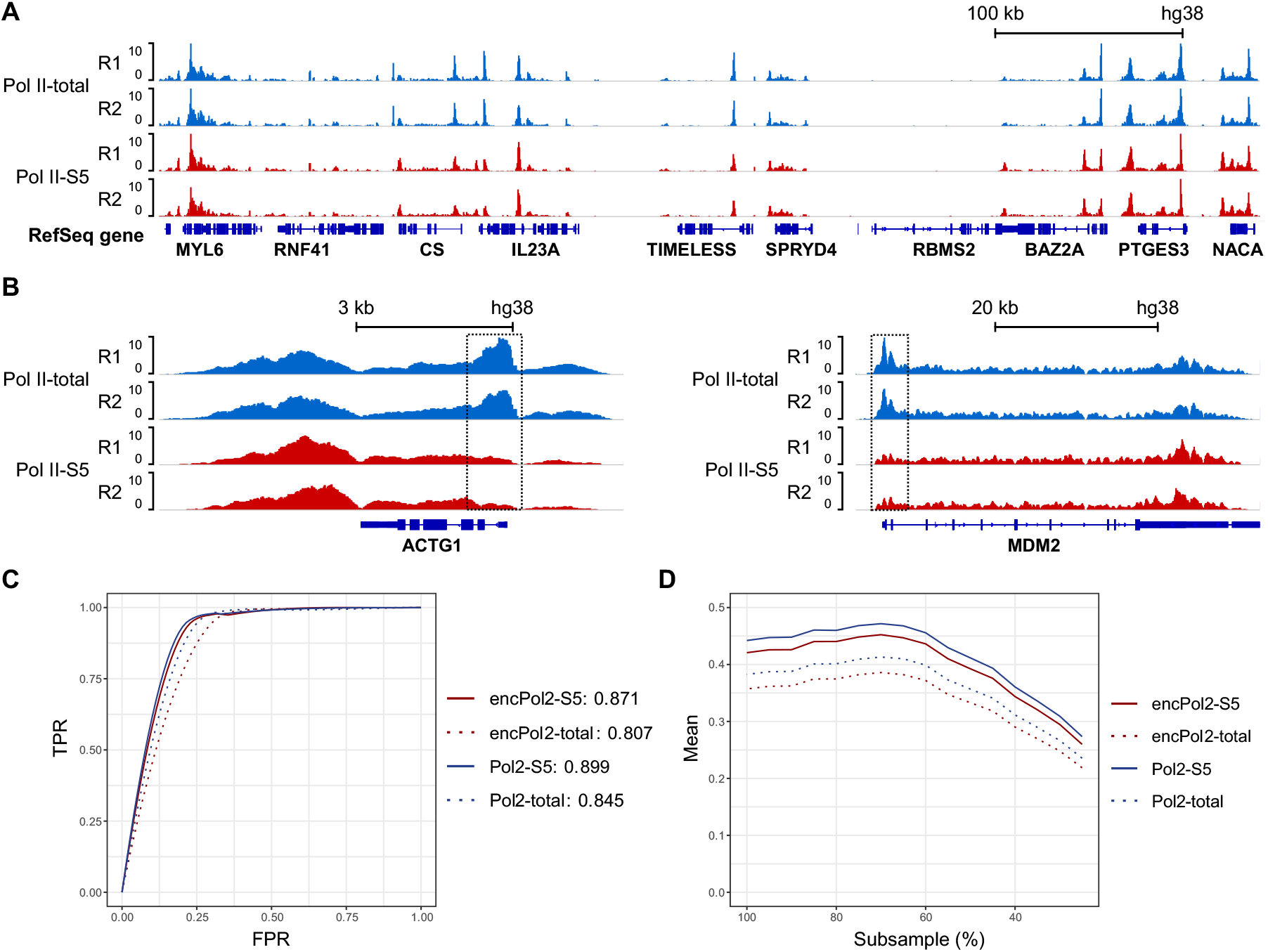
Comparison of Pol II-total and Pol II-S5 bindings in GM12878 cells. (A) Normalized Pol II-S5 and Pol II-total signal in GM 12878 cells (50,000 cells per assay). (B) Normalized Pol II signal over genes ACTG1 and MDM2, showing the difference in Pol II-total and Pol II-S5 binding. Pol II-total includes Pol II that is not phosphorylated. These non-active Pol II molecules tend to pause at promoter regions, while phosphorylated and active Pol II (S5 being the majority) binds to the entire gene body. (C) Receiver operating characteristic (ROC) curves using ENCODE (SRX100400 and SRX100530) and nMOWChIP-seq Pol II-total and Pol II S5 datasets predicting EGR1 binding peaks in GM12878 cells. (D) Matthew’s correlation coefficient (MCC) plots of ENCODE and nMOWChIP-seq datasets being subsampled at different levels to predict EGR1 binding peaks in GM12878 cells.

To determine whether Pol II-total or Pol II-S5 was more effective in predicting TF binding sites, we analyzed the overlap of either Pol II-total peaks or Pol II-S5 peaks with EGR1 peaks at varying peak-calling thresholds, shown as receiver operating characteristic (ROC) curves that display the true positive rate (TPR) versus the false positive rate (FPR) (Fig. 2c). To quantify the predictive quality, we calculated the area under the curve (AUC) for each of the ROC curves using our EGR1 binding data as the gold standard. We found that Pol II-S5 profile had a higher predictive value than Pol II-total (0.899 vs 0.845 using nMOWChIP-seq data, and 0.871 vs 0.807 using ENCODE data). We also tested if Pol II-S5 was more robust than Pol II-total with samples of lower quality (Fig. 2D). For this, we subsampled our EGR1 data from 25% to 95% of the original reads, with 5 replicates at each sampling. We then calculated the average Matthew’s correlation coefficient (MCC), a robust single value quantifier of classifier quality, of the five replicates at each subsampling percentage. Pol II-S5 outperformed Pol II-total, both in our data and in ENCODE’s, regardless of TF data quality.

### Differential RNA Pol II binding in prefrontal cortex and cerebellum of mouse brain

We applied nMOWChIP-seq to profile Pol II binding in mouse PFC and cerebellum. PFC have roles in cognitive functions, decision-making and short-term memory,^38,39^ while cerebellum controls motor functions and coordination.^40^ We reasoned that the functional difference between the two regions should reflect on the binding of Pol II and TFs that have recognized roles in the brain. There have not been published ChIP-seq data confirming this.

We mapped Pol II-total binding in B6 mouse brain using nuclei extracted from PFC and cerebellum (Fig. 3). The genome-wide Pol II profiles were substantially different between cerebellum and PFC with Pearson’s correlation being 0.67, compared to an average of 0.96 between technical replicates (Fig. 3A and 3B). DiffBind analysis identified 3021 peaks with higher levels of Pol II binding in PFC than in cerebellum, and 1197 peaks having higher binding in cerebellum (fold change > 2, p < 10^−5^). Gene ontology (GO) analysis of these regions showed that the regions with high Pol II binding intensity in PFC were enriched in terms associated with memory, learning and anxiety-related response (Fig. 3C). Synaptic plasticity, one of the fundamentals of learning and memory, was also enriched along with its key components: long term potentiation and depression (Figure 3C).^41^ In contrast, the genomic regions with Pol II binding higher in cerebellum were enriched in terms that were specific to cerebellum (e.g. cerebellum morphology and development) (Fig. 3C).

**Fig 3.**
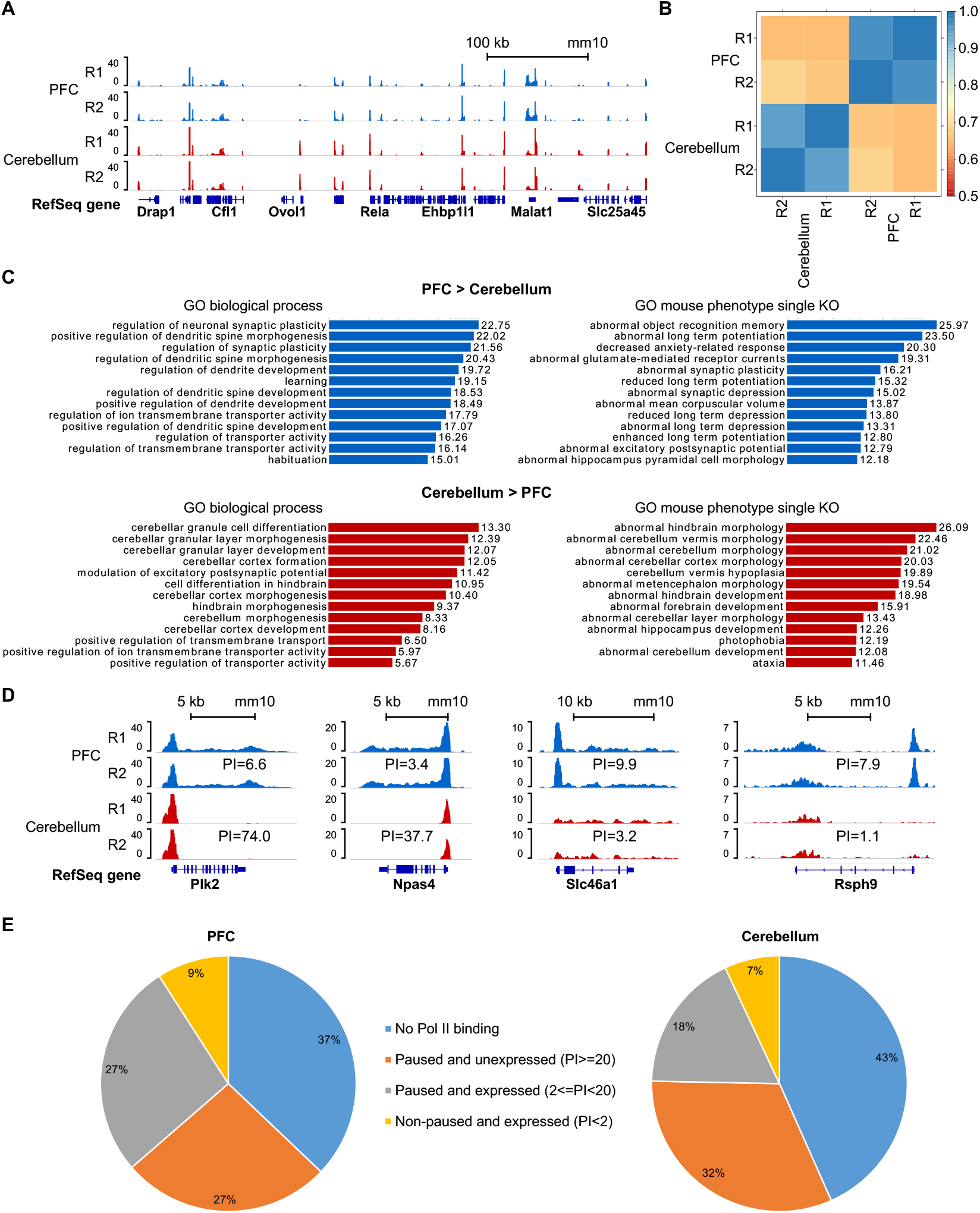
Differential RNA polymerase II binding in prefrontal cortex and cerebellum of mouse brain. (A) Normalized Pol II-total signals generated using nuclei isolated from mouse prefrontal cortex (PFC) and cerebellum (50,000 nuclei per assay). (B) Pearson’s correlations of affinity score calculated from consensus MACS2 peaks and uniquely mapped reads using DiffBind, among Pol II data on PFC and cerebellum. (C) GO biological processes and mouse phenotypes (single knockout) associated with regions having higher Pol II binding levels in either PFC or cerebellum (-log10 binomial p value, fold change > 2, p < 10^−5^). (D) Normalized Pol II-total signals at genes that have significantly different pausing indexes between PFC and cerebellum. (E) Distribution of Pol II pausing index in PFC and cerebellum.

We also examined the Pol II pausing index for the Pol II-bound genes in PFC and cerebellum. Pol II pausing index (PI) refers to the ratio of Pol II read density between promoter proximal region (−30 to +300 bp of TSS) and gene body (+300 bp of TSS to TES), with higher PI indicating more Pol II binding near TSS^2,42^. Pol II-bound genes can be divided into three categories based on the PI value: non-paused and expressed (PI<2), paused and expressed (2<PI<20), and paused and unexpressed (PI>20).^2^ For example, Plk2, which is a gene known to participate in rodent brain development and cell proliferation,^43^ and Npas4, which is involved in regulating reward-related learning and memory,^44^ both showed dramatically higher PI values in cerebellum than in PFC (Fig. 3D). The data revealed that they were actively expressed in PFC but paused without expression in cerebellum. In contrast, Slc46a1 and Rsph9 showed higher PI values in PFC than cerebellum. While both were being expressed in the two tissues, there was significant pausing of Pol II at TSS in PFC. This suggests that PFC had more potential in transcribing these genes at a short notice, such as Slc46a1, which encodes a facilitative carrier for folate.^45^ Furthermore, we analyzed the distribution of the PI in PFC and cerebellum (Fig. 3E). 36% of genes were being actively expressed in PFC (PI<20), compared to only 25% in cerebellum. Mouse brain displays more pausing and less active transcription compared to GM12878 (52% actively expressed genes, Supplementary Fig. S5). This was also within our expectation because GM12878 cell line was maintained in log phase and actively dividing, unlike brain cells in adult mice. We identified Pol II-bound genes that displayed significantly different pausing indexes between PFC and cerebellum (fold change > 3, minimal read density > 0.02 read/bp) (Supplementary Table S2). Among these, Igfbp6 is involved in myelin formation during central nervous system (CNS) development,^46^ and Slc1a2 encodes excitatory amino acid transporter 2, responsible for reuptake of 90% glutamate in CNS.^47^ Shank3 belongs to the Shank gene family that plays a role in synapse formation,^48^ and Flrt2 encodes a member of the FLRT protein family that is shown to regulate signaling during mouse development.^49^

### Differential EGR1 and MEF2c binding in prefrontal cortex and cerebellum of mouse brain

We also examined EGR1 and MEF2C (myocyte enhancer factor-2 C) binding using nuclei extracted from mouse PFC and cerebellum. Chromatin from 100,000 nuclei were used in each assay, yielding high quality data that revealed differential binding between the two regions of brain (Figure 4A). We picked several genes as examples (Fig. 4A). Kalrn (Kalirin) plays important roles in nerve growth^50^. Higher EGR1/MEF2C activity in PFC was observed on Gria1(Glutamate receptor 1) which is involved in synaptic transmission.^51^ Cacna1a is involved in movement disorder^52^ and expression of Nfix can influence neural stem cell differentiation.^53^ Zic1 and Zic4 belong to the family of Zinc finger of the cerebellum (ZIC) protein family,^54^ whose loss of function can lead to Dandy-Walker malformation and incomplete cerebellar vermis^55^. Higher EGR1 binding on these four genes were seen in cerebellum than in PFC. A large fraction of EGR1 and MEF2C peaks (68-89%) appear to overlap with Pol II peaks, in both cerebellum and PFC (Supplementary Fig. S6). We observed correlated EGR1 and MEF2C profiles, with an average of Pearson’s correlation of 0.74 between the two TFs in PFC and 0.84 in cerebellum (Fig. 4B). Such correlations were much higher than the one observed in GM12878 cells (r ~0.52). On the other hand, EGR1 and MEF2C present very different profiles between PFC and cerebellum, with the average correlation of 0.51 between the two brain regions for both TFs. We further examined the binding difference between PFC and cerebellum for these two TFs. Their binding sites were much more heavily situated at promoters in PFC (79%) than in cerebellum (60%) (Fig. 4C). We further analyzed the data using DiffBind to identify regions with significantly different level of EGR1/MEF2C binding between PFC and cerebellum.^56^ In total, DiffBind identified 1026 peaks with higher EGR1 binding in cerebellum than in PFC, and 1563 peaks with higher MEF2C binding in cerebellum than in PFC (fold change > 2, p < 10^−5^). These regions were further analyzed for GO term enrichment analysis using GREAT and linked to walking behavior, motor coordination, and cerebellum morphology and development in the case of EGR1, cerebellar development and limb coordination in the case of MEF2C (Figure 4D). In contrast, very few differential peaks (12 for EGR1 and 1 for MEF2C) were found to have higher binding signal in PFC than in cerebellum and no GO terms were found.

**Fig 4.**
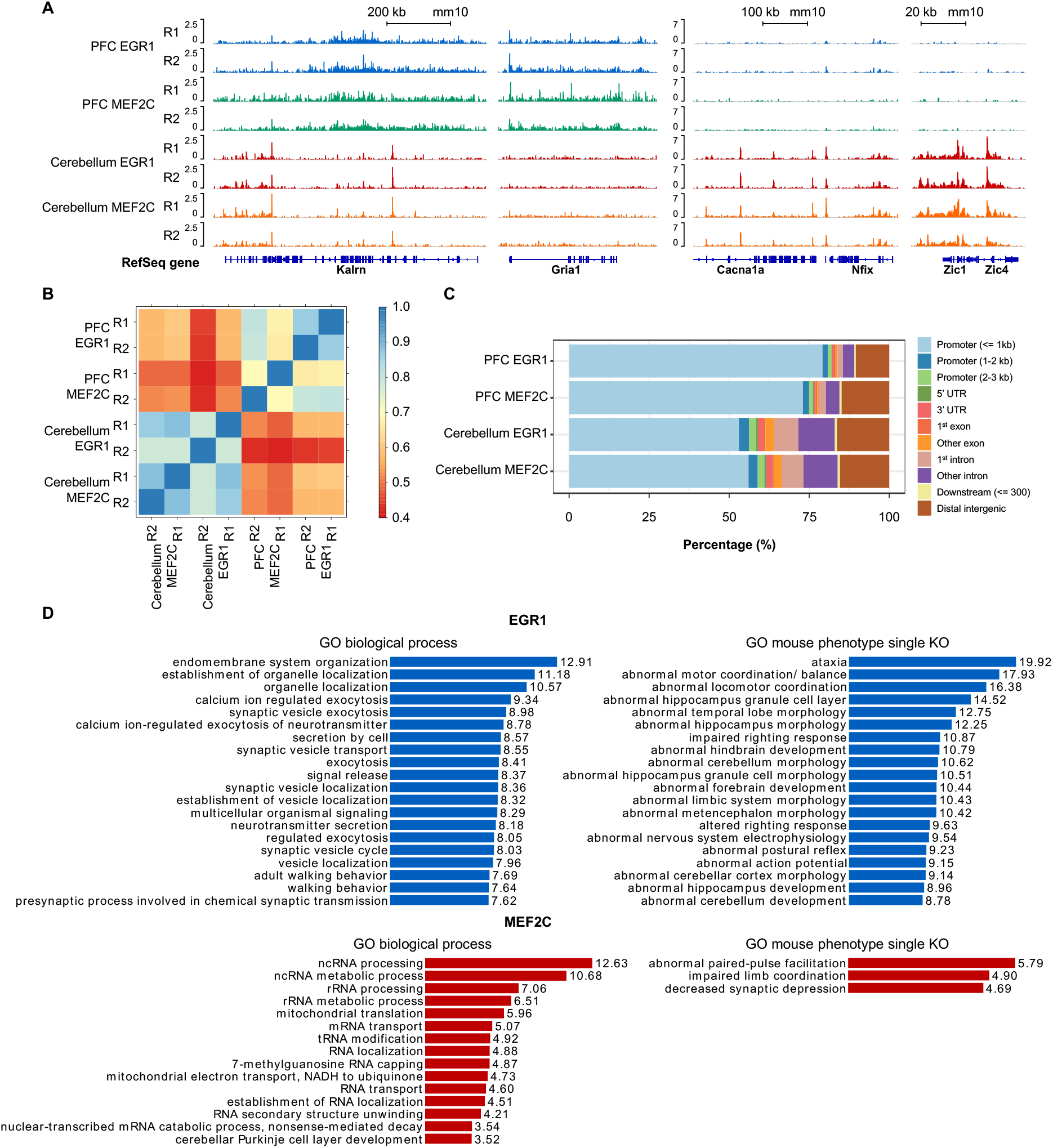
Differential transcription factor binding in prefrontal cortex and cerebellum of mouse brain. (A) Normalized EGR1 and MEF2C signals at genes identified by DiffBind to have significantly different binding between PFC and cerebellum (fold change > 2, p < 10^−5^). (B) Pearson’s correlations of affinity score calculated from consensus MACS2 peaks and uniquely mapped reads using DiffBind, among TF data on PFC and cerebellum. (C) Distribution of TF binding peaks (p < 10^−5^) in various genomic regions. Each peak set is the overlapping peaks from 2 replicates of the same sample. (D) GO biological process and mouse phenotype (single knockout) terms associated with regions having higher EGR1 and MEF2C binding levels in cerebellum than PFC (-log10 binomial p value, fold change > 2, p < 10^−5^).

## Discussion

ChIP-seq profiling of proteins bound to the genome is generally much more challenging than that of modified histones. Histone ChIP-seq can be conducted with crosslinking and sonication or under native ChIP condition. Due to the robust interaction between histones and genome, native ChIP-seq for histone modifications can have very high efficiency. In contrast, previous ChIP-seq of protein bindings often involves crosslinking and sonication that immobilizes the protein to the interacting DNA sequence before breaking chromatin into fragments. Crosslinking is often considered necessary when TF, Pol II and enzyme interaction with the genome is studied, due to perceived needs and benefits for preservation of such interactions by crosslinking. However, crosslinking and sonication potentially cause epitope masking^57^ and damage, respectively, and both affect antibody-antigen interaction critically involved in ChIP assays. Furthermore, crosslinking may also create artifact peaks at highly transcribed regions due to protein-protein crosslinking^58^.

In this work, by conducting nMOWChIP-seq without crosslinking and sonication, we demonstrate a low-input technology that works with as few as 1,000-50,000 cells per assay for profiling a wide range of protein-genome interactions. We show that nMOWChIP-seq method effectively preserves the interaction between Pol II/TFs/enzyme and the genome. In our approach, the required number of cells depends on the robustness of the protein binding to the genome under native ChIP conditions, the number of binding sites, and the quality of the antibody. Compared to the state-of-the-art ChIP-seq data taken using millions of cells per assay, our datasets generally show very high signal-to-noise ratio and low background, which are characterized by high FRiP values. Our method is also comparable to CUT&RUN^24,27^ in terms of data quality.

Compared to histone modification data, ChIP-seq data on Pol II, TFs, enzyme in tissues are very scarce. We applied nMOWChIP-seq to profile Pol II, and key TFs EGR1 and MEF2C in a brain-region-specific manner in cerebellum and prefrontal cortex of mouse brain. We found that Pol II and TF profiles are highly characteristic of the brain regions. The Pol II binding profiles had 4,218 differential peaks between cerebellum and PFC, while EGR1/MEF2C profiles had >1,000 differential peaks between the two brain regions. The peaks with their intensity high in PFC and low in cerebellum were highly enriched in functions including cognition, learning or memory, while the peaks that were high in cerebellum and low in PFC were enriched in GO terms including walking behavior, motor coordination and cerebellar cortex formation. The fact that the binding profiles of these functional molecules are highly characteristic of the functions of various brain regions indicate that Pol II and the key TFs are critically involved in the molecular dynamics associated with the spatial configuration of brain functions. Our results suggest the possibility of deciphering genome-wide molecular binding profiles to establish the neuroanatomical origin of brain tissues for the first time.

## Methods

### Cell culture

GM12878 cells were obtained from Coriell Institute for Medical Research. Cells were cultured in RPMI-1640 medium (30-2001, ATCC) with 15% fetal bovine serum (16000-044, Gibco) and 1% pen-strep (Invitrogen) at 37°C, 5% CO_2_. Cells were subcultured every 3 d to maintain exponential growth.

### Mouse strain and brain dissection

C57BL/6J mice were purchased from Jackson Laboratory and maintained in the animal facility with 12-h light/12-h dark cycles and food and water ad libitum. 8-week old male mice were sacrificed by compressed CO_2_ followed by cervical dislocation. Mouse brains were rapidly dissected, frozen on dry ice and stored at −80°C. This study was approved by the Institutional Animal Care and Use Committee (IACUC) at Virginia Tech.

### Nuclei isolation from brain tissues

A mouse brain was put on ice and PFC and cerebellum were dissected for nuclei isolation. The following steps are performed on ice and centrifugation performed at 4°C. Tissue was placed in 3 ml of ice-cold nuclei extraction buffer [0.32 M sucrose, 5 mM CaCl_2_, 3 mM Mg(Ac)_2_, 0.1 mM EDTA, 10 mM tris-HCl, and 0.1% Triton X-100, with 30 μl of PIC (P8340, Sigma-Aldrich), 3 μl of 100 mM PMSF, and 3 μl of 1 M dithiothreitol added before use]. Tissue was homogenized in the grinder set (D9063, Sigma-Aldrich) by slowly douncing 15 times with pestle A and 25 times with pestle B. Homogenate was filtered through a 40 μm cell strainer into a 15 ml tube and centrifuged at 1000*g* for 10 min. The supernatant was removed and the pellet was resuspended in 500 μl nuclei extraction buffer and transferred to a 1.5 ml tube. 750 μl of 50% iodixanol, 7.5 μl of PIC, 0.75 μl of 100 mM PMSF and 0.75 μl of 1M dithiothreitol were added and mixed by pipetting. The mixture was centrifuged at 10,000*g* for 20 min and the supernatant was removed. If mixed nuclei (without separation of neurons and glia) were used for MOWChIP directly, the nuclei pellet was resuspended in 200 μl of Dulbecco’s phosphate-buffered saline (DPBS). If nuclei labeling and FACS sorting were conducted, 500 μl of 2% normal goat serum (50062Z, Life Technologies) in DPBS was added to the nuclei pellet and incubated for 10 min before resuspending. Anti-NeuN antibody conjugated with Alexa 488 (MAB377X, EMD Millipore) was diluted with DPBS to 2 ng/μl. 8 μl of anti-NeuN was added to each 500 μl of nuclei suspension and incubated at 4°C for 1 h on a rotator. The labeled nuclei were then sorted using FACS (BD FACSAria, BD Biosciences). 8 μl of non-labeled nuclei were saved as unstained control prior to addition of anti-NeuN antobody. The concentration of nuclei suspension after FACS was typically low (~1.2×10^5^/ml). The nuclei were re-concentrated by adding 200 μl of 1.8 M sucrose, 5 μl of 1M CaCl_2_ and 3 μl of 1M Mg(Ac)_2_ to 1 ml of the nuclei suspension. The mixture was incubated on ice for 15 min and centrifuged at 1800*g* for 15 min. Supernatant was removed and the nuclei pellet was resuspended in DPBS.

### MNase digestion of chromatin

This protocol is scalable in volume and can digest cell/nuclei suspension with concentration up to 4×10^6^/ml. Our experiments usually started with 4×10^5^ cells/nuclei suspended in 100 μl of DPBS. 1 μl of PIC, 1 μl of 100 mM PMSF and 100 μl lysis buffer [4% Triton X-100, 100 mM tris-HCl, 100 mM NaCl, and 30 mM MgCl_2_] were added, mixed by vortexing and incubated at room temperature for 10 min. 10 μl of 100 mM CaCl_2_ and 2.5 μl 100U MNase (88216, Thermo Fisher Scientific) were added, mixed by vortexing and incubated at room temperature for 10 min. 22 μl of 0.5 M EDTA was then added, mixed by vortexing and incubated on ice for 10 min. The solution was centrifuged at 16,100*g* for 5 min at 4°C. Supernatant containing fragmented chromatin was collected into a new 1.5 ml tube and placed on ice for use.

### Preparation of immunoprecipitation (IP) beads

5 μl of protein A Dynabeads (10001D, Invitrogen) were used in each MOWChIP assay. The beads were washed twice with IP buffer (20 mM Tris-HCl, pH 8.0, 140 mM NaCl, 1 mM EDTA, 0.5 mM EGTA, 0.1% (w/v) sodium deoxycholate, 0.1% SDS, 1% (v/v) Triton X-100) and resuspended in 150 μl of IP buffer. 1 μg of Pol II or TF antibody was added into the bead suspension for each assay using 10^5^ cells/nuclei, while 0.5 μg of antibody was added for each assay using 5×10^4^ or fewer cells/nuclei. We used the following antibodies in this work: anti-Pol II-total (ab817, lot GR3271868-2, Abcam), anti-Pol II-S5 (ab5131, lot GR3202335-5, Abcam), anti-EGR1 (sc-101033, lot A1618, Santa Cruz), anti-MEF2C (sc-365862, lot B0818, Santa Cruz), anti-HDAC2 (ab124974, lot GR97402-7, Santa Cruz). The suspension was incubated on a rotator at 4°C for 2 h. The beads were then washed with IP buffer twice and resuspended in 5 μl of IP buffer for loading into the device chamber.

### MOWChIP-seq

The microfluidics-based MOWChIP-seq process, including microfluidic device fabrication and operation, was conducted following our published protocol^14^.

### Purification of ChIP DNA

After MOWChIP, IP beads were rinsed once with IP buffer and resuspended in 200 μl of DNA elution buffer (10 mM Tris-HCl, 50 mM NaCl, 10 mM EDTA, and 0.03% SDS). 2 μl of 20 mg/ml proteinase K was added and the suspension was incubated at 65°C for 1 h. DNA was then extracted and purified by phenol-chloroform extraction and ethanol precipitation. DNA pellet was resuspended in 8 μl of low EDTA TE buffer. Input DNA was purified with the same process, by digesting chromatin solution directly using proteinase K, followed by the same extraction and purification process.

### Library preparation, quantification and sequencing

Libraries were constructed using Accel-NGS 2S Plus DNA Library kit (Swift Biosciences) following the manufacturer’s instructions. 1 × EvaGreen dye (Biotium) was added to the amplification mixture to monitor the PCR amplification. Library was eluted to 7 μl of low EDTA TE buffer. Library fragment size was examined using TapeStation (Agilent) and the concentration was quantified with KAPA Library Quantification kit (Kapa Biosystems). We also examined the enrichment of the libraries using qPCR and primers listed in Supplementary Table S3. Libraries were pooled for sequencing by Illumina HiSeq 4000 SR50 mode.

### ChIP-seq data analysis

Raw sequencing FASTQ files were trimmed using Trim Galore! with default settings. Reads that passed the quality check were mapped to reference genomes hg38 (human) or mm10 (mouse) with Bowtie.^59^ Uniquely mapped reads were filtered for known blacklisted genome regions using Samtools^60^ and bedtools^61^ to remove ChIP-seq artifacts. The reference genome was then divided into 100-bp bins and ChIP-seq signal for each bin was counted for both ChIP and input samples. Normalized ChIP-seq signal for each bin was calculated using the following equation:

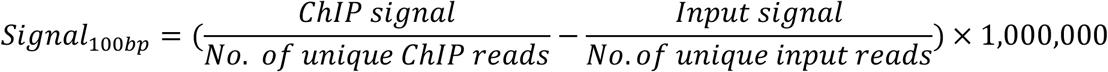

Both ChIP and input reads were then extended by 100 bp on either ends to compute normalized signal for each bin in the same manner, and visualized as tracks in genome browser IGV.^62^ Peak calling was conducted using MACS2 (default settings). Differential binding analysis was carried out using DiffBind^56^ with default settings. Selected genomic regions of interest were further analyzed for gene ontology terms using GREAT.^63^ Pearson’s correlation between datasets was calculated via DiffBind, using *dba.count* command with default parameters to process MACS2 peaks and uniquely mapped reads and calculate consensus peaks and affinity score.

ROC curves were created by calculating the true positive rate (TPR) and false positive rate (FPR) at 100 different peak-calling thresholds from 0.99 and 10^−15^. Thresholds were applied to both the EGR1 and Pol II sets, and AUC was calculated with the R package ROC. Subsampling of EGR1 bam files to between 25% to 95% of original reads was performed using Samtools. Five separate, but consistent, seeds were used at each subsampling percentage. Matthew’s correlation coefficient (MCC) was calculated in R (mltools) at each percentage and each seed using a MACS2 q-value cutoff of 0.05 for both EGR1 and Pol II peaks.

## Data Availability

The ChIP-seq data sets are deposited in the Gene Expression Omnibus (GEO) repository with the following accession number GSE172224. https://www.ncbi.nlm.nih.gov/geo/query/acc.cgi?acc=GSE172224

## Acknowledgements

This work was supported by US National Institutes of Health (NIH) grants R33 CA214176 (C.L.), R01EB017235 (C.L.), R01 CA243249 (C.L.), P30 CA012197 (C.L.), R01NS094574 (H.X.), R21MH120498 (H.X.), and a seed grant from Virginia Tech Institute for Critical Technology and Applied Science (C.L.).

## Author Contribution

C.L. designed and supervised the study. Z.L. conducted all nMOWChIP-seq experiments and analyzed the data. L.B.N., Y.Z., C.D., Q. Z., B.Z., Z.Z., M.S. helped with data analysis and experiments. A.M. and H.X. helped with experiments on the TFs. Z.L., L.B.N. and C.L. wrote the manuscript. All authors proofread the manuscript and provided feedback.

