## Supplementary Figures and Tables for "Native MOWChIP-seq: Genome-wide profiles of key protein bindings reveal functional differences among various brain regions"

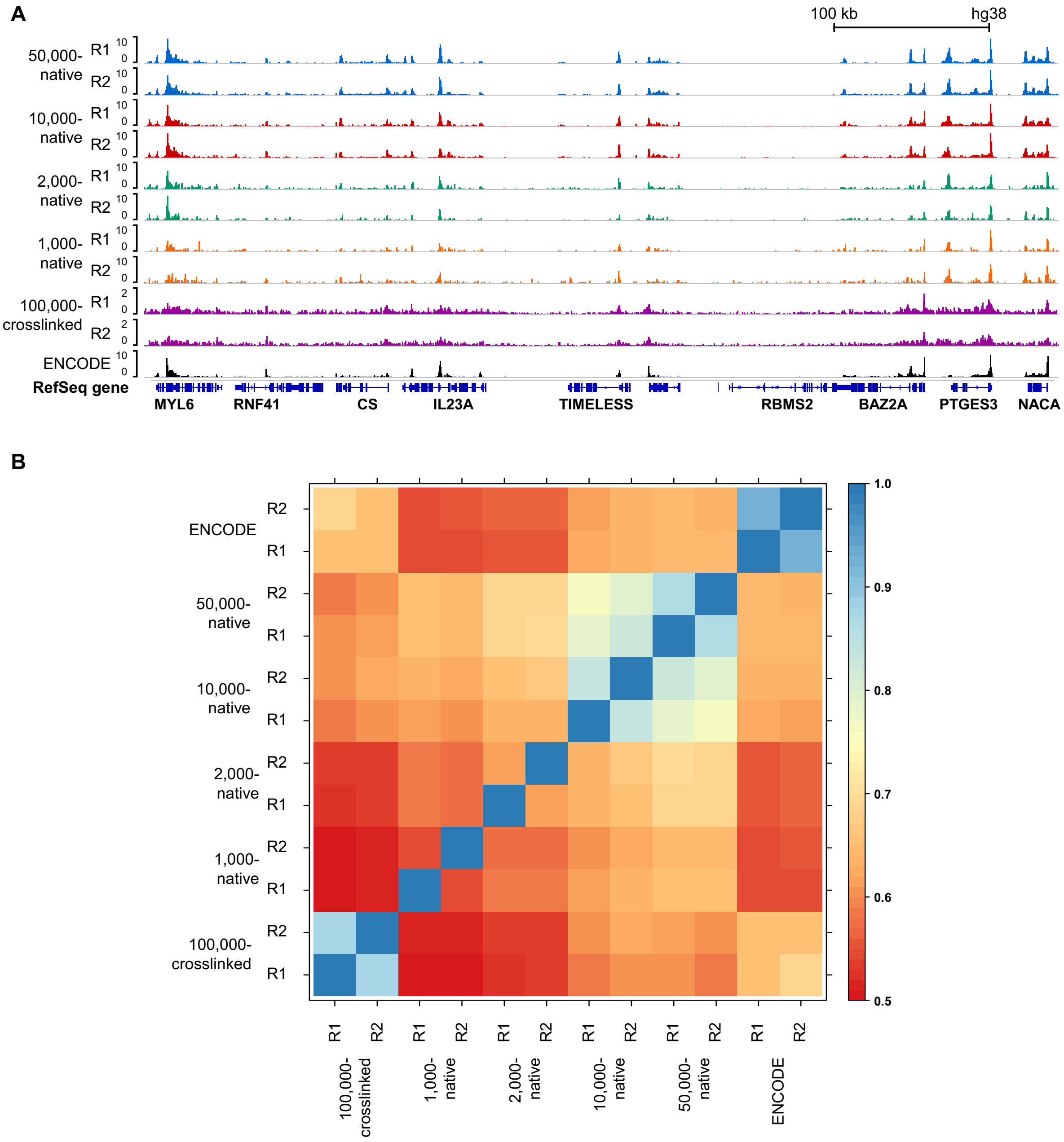

**Supplementary Fig S1. Comparison of MNase-based native MOWChIP-seq with crosslinked MOWChIP-seq.** (A) Normalized Pol II (S5) signal of GM12878 using MOWChIP-seq, with either native (1,000-50,000 cells per assay) or crosslinked (100,000 cells per assay) chromatin. (B) Pearson's correlation matrix of native and crosslinked MOWChIP-seq data and ENCODE data (SRX100530), computed using DiffBind affinity score method.

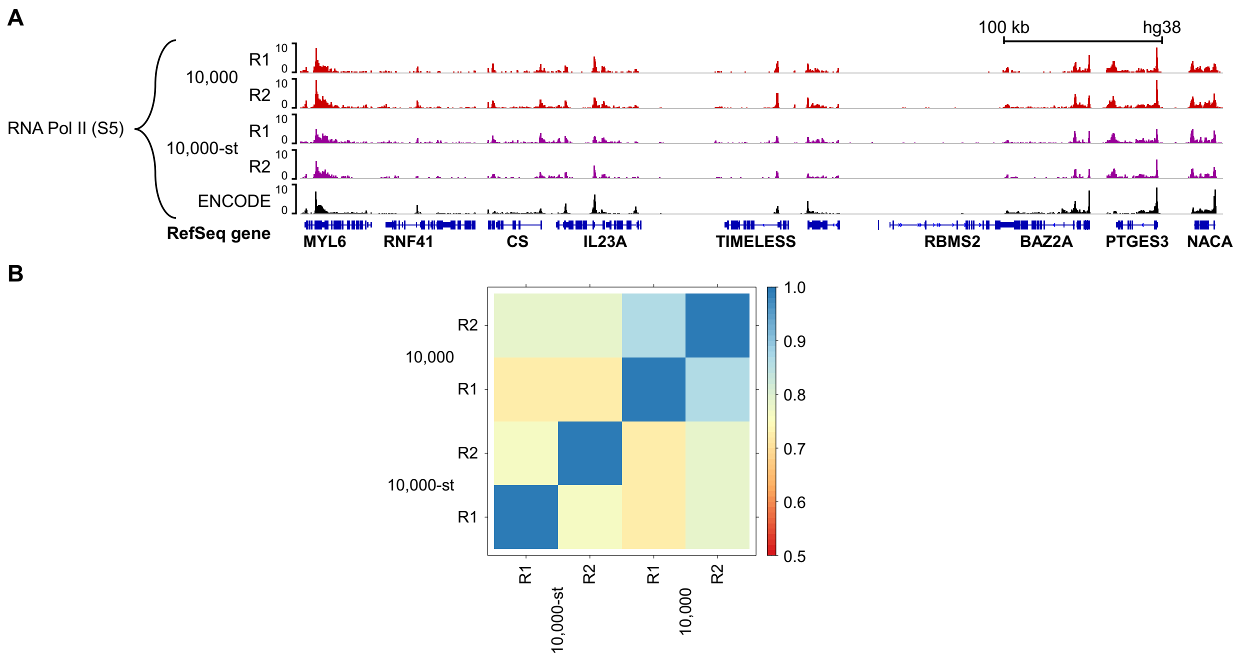

**Supplementary Fig S2. Storage at -80 °C does not change RNA pol II binding profile.** (A) Normalized Pol II-S5 signal of GM12878 cells. “10,000-st” denotes chromatin samples (each containing chromatin fragments from 10,000 cells) that were stored at -80 °C for 2 d and then used for nMOWChIP-seq. “10,000” refers to samples processed without such storage. In contrast, we found that freezing of chromatin samples at -80 °C substantially varies TF binding profiles. (B) Pearson’s correlation matrix of Pol II-S5 nMOWChIP-seq data of fresh and -80 °C stored GM12878 chromatin, computed using DiffBind affinity score method.

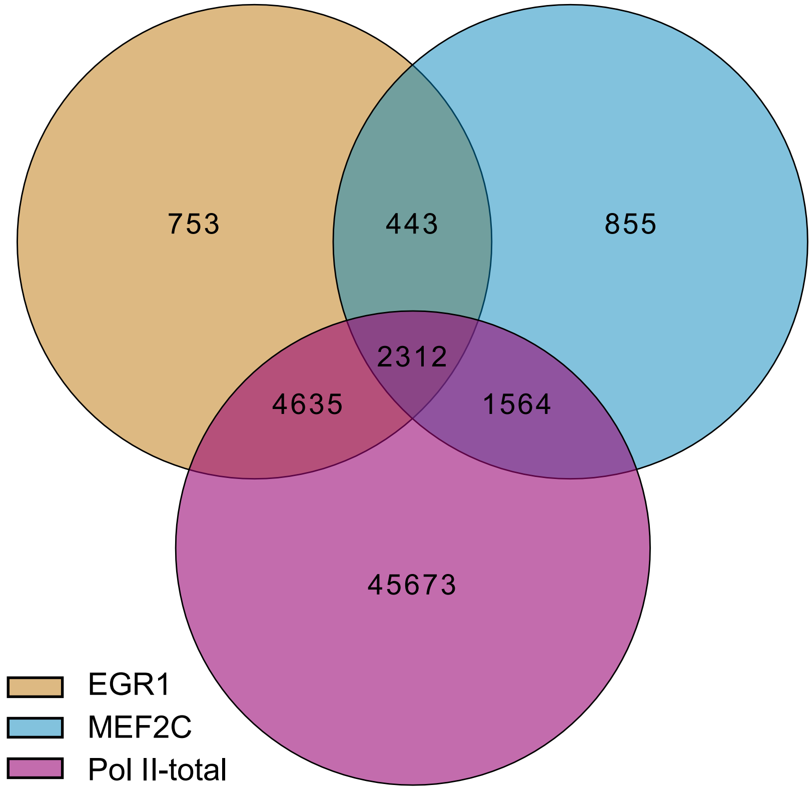

**Supplementary Fig S3. Overlap among Pol2-total, EGR1 and MEF2C peaks of GM12878 cells.** Peaks were identified by MACS2 using default settings. Peaks from both replicates that overlap more than 50% of their length are merged into a common peak using bedtools. The overlap of the common peaks was plotted using Intervene.

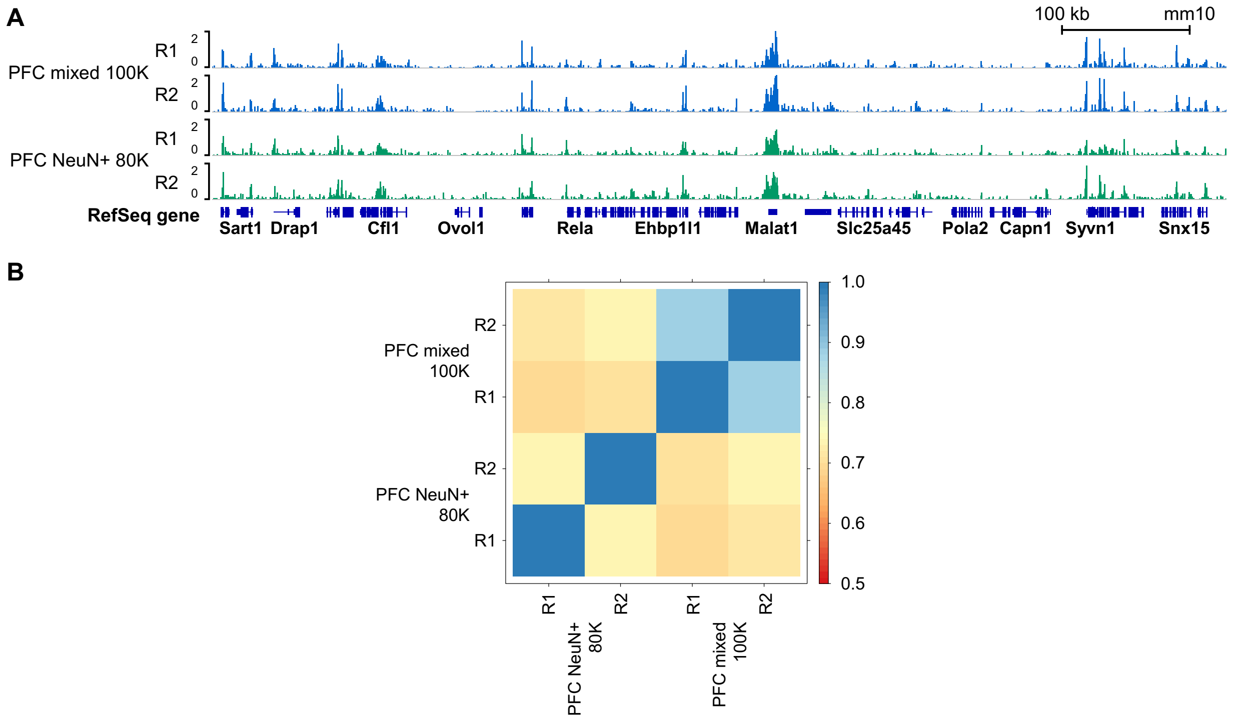

**Supplementary Fig S4. HDAC2 binding in mouse PFC samples profiled by nMOWChIP-seq.** (A) Normalized HDAC2 signals generated using nuclei from mouse PFC. 100,000 unsorted nuclei from PFC, and 80,000 of FACS-sorted NeuN+ (neuronal) nuclei were used per assay in these tests. (B) Pearson's correlation matrix on HDAC2 mouse PFC data, computed using DiffBind affinity score method.

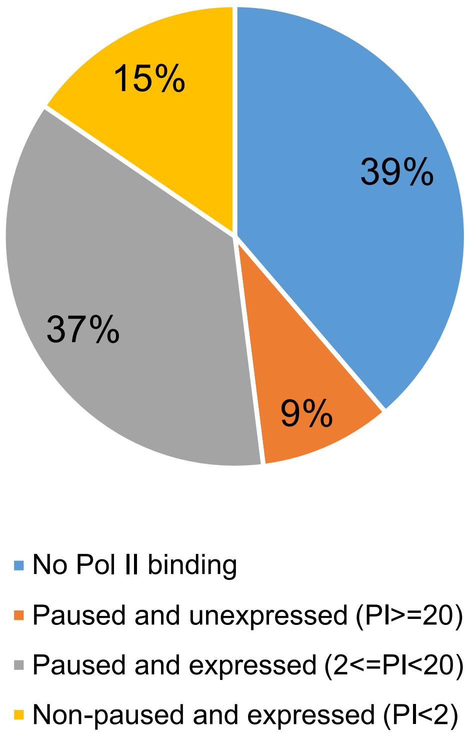

**Supplementary Fig S5. Distribution of pausing index (PI) for genes in GM12878 cells based on Pol II-total nMOWChIP-seq data.**

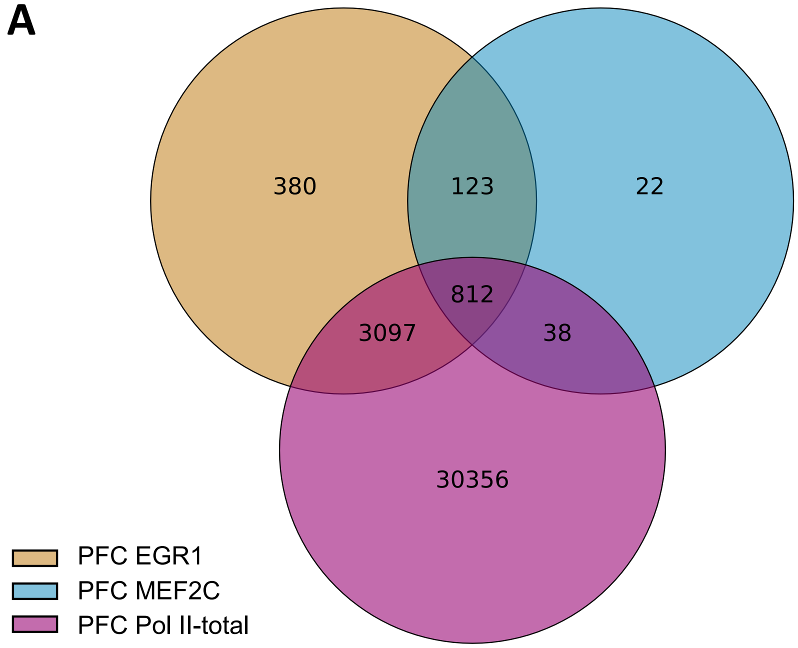

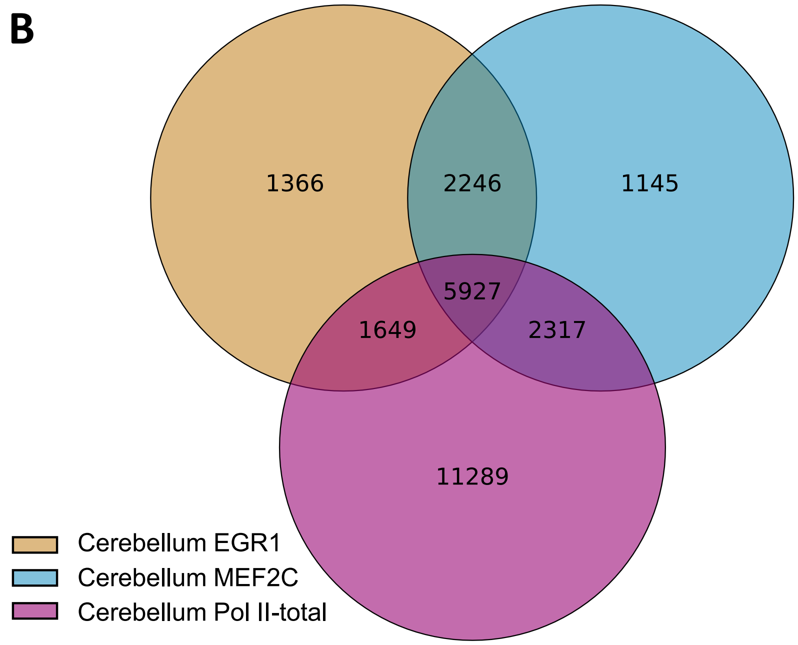

**Supplementary Fig S6. Overlap among Pol II, EGR1, and MEF2C binding peaks in mouse PFC (A) and cerebellum (B) data.** Peaks were identified by MACS2 using default settings. Peaks from both replicates that overlap more than 50% of their length are merged into a common peak using bedtools. The overlap of the common peaks was plotted using Intervene.

**Supplementary Table S1. Summary of nMOWChIP-seq data**

| File | Mapped reads | Mapping Rate | Unique reads | Unique rate | MACS2 | FRIP |
|---|---|---|---|---|---|---|
| GM12878_Pol2-total_50K_1 | 16432424 | 98% | 16204517 | 99% | 133015 | 57.1% |
| GM12878_Pol2-total_50K_2 | 20219467 | 98% | 19820603 | 98% | 129219 | 55.5% |
| GM12878_Pol2-S5_50K_1 | 16006273 | 98% | 15570715 | 97% | 142339 | 52.4% |
| GM12878_Pol2-S5_50K_2 | 15203135 | 98% | 14377380 | 95% | 130669 | 47.3% |
| GM12878_Pol2-S5_10K_1 | 12271356 | 96% | 12061753 | 98% | 114811 | 41.3% |
| GM12878_Pol2-S5_10K_2 | 21780356 | 96% | 21550914 | 99% | 140695 | 48.0% |
| GM12878_Pol2-S5_10K_st_1 | 9646207 | 95% | 9354458 | 97% | 82320 | 27.8% |
| GM12878_Pol2-S5_10K_st_2 | 13775137 | 96% | 13338787 | 97% | 79800 | 24.9% |
| GM12878_Pol2-S5_2K_1 | 14243331 | 90% | 13432201 | 94% | 59713 | 17.7% |
| GM12878_Pol2-S5_2K_2 | 15242929 | 87% | 14400694 | 94% | 70439 | 22.0% |
| GM12878_Pol2-S5_1K_1 | 15572762 | 87% | 14923018 | 96% | 33627 | 10.7% |
| GM12878_Pol2-S5_1K_2 | 9194036 | 86% | 8777057 | 95% | 19820 | 8.1% |
| GM12878_Pol2-S5_100K_cl_1 | 31745569 | 98% | 31406005 | 99% | 53817 | 8.8% |
| GM12878_Pol2-S5_100K_cl_2 | 31111394 | 98% | 30545134 | 98% | 17561 | 2.7% |
| GM12878_EGR1_100K_1 | 14323523 | 96% | 12199195 | 85% | 16614 | 8.3% |
| GM12878_EGR1_100K_2 | 17458334 | 96% | 14961426 | 86% | 15584 | 7.9% |
| GM12878_EGR1_50K_1 | 14910367 | 95% | 12800282 | 86% | 16465 | 8.6% |
| GM12878_EGR1_50K_2 | 16669698 | 95% | 14480504 | 87% | 15159 | 8.2% |
| GM12878_EGR1_10K_1 | 9072849 | 84% | 7725182 | 85% | 2626 | 3.8% |
| GM12878_EGR1_10K_2 | 12461029 | 94% | 10785293 | 87% | 2781 | 2.6% |
| GM12878_EGR1_5K_1 | 10330104 | 92% | 8383719 | 81% | 8245 | 5.0% |
| GM12878_EGR1_5K_2 | 12660454 | 93% | 11175451 | 88% | 1423 | 1.9% |
| GM12878_MEF2C_100K_1 | 24369737 | 76% | 21133894 | 87% | 14716 | 4.5% |
| GM12878_MEF2C_100K_2 | 24642860 | 85% | 20705206 | 84% | 19884 | 6.3% |
| GM12878_HDAC2_100K_1 | 21068312 | 84% | 18801502 | 89% | 11221 | 3.8% |
| GM12878_HDAC2_100K_2 | 19862691 | 85% | 18127467 | 91% | 5832 | 2.3% |
| GM12878_HDAC2_50K_1 | 8767036 | 93% | 8093468 | 92% | 1204 | 1.3% |
| GM12878_HDAC2_50K_2 | 12215492 | 93% | 11358639 | 93% | 1584 | 1.1% |
| PFC_Pol2-total_50K_1 | 13064208 | 97% | 12803650 | 98% | 66123 | 48.6% |
| PFC_Pol2-total_50K_2 | 16395371 | 99% | 16251965 | 99% | 80138 | 55.4% |
| PFC_Pol2-total_10K_1 | 8574476 | 95% | 7748982 | 90% | 24423 | 26.5% |
| PFC_Pol2-total_10K_2 | 13007858 | 93% | 12338679 | 95% | 53291 | 46.4% |
| Cerebellum_Pol2-total_50K_1 | 10617272 | 99% | 10444570 | 98% | 43387 | 52.0% |
| Cerebellum_Pol2-total_50K_2 | 9805738 | 98% | 9553592 | 97% | 36762 | 48.5% |
| Cerebellum_Pol2-total_10K_1 | 15483210 | 97% | 15200342 | 98% | 54534 | 67.1% |
| Cerebellum_Pol2-total_10K_2 | 14308273 | 98% | 14139656 | 99% | 54278 | 69.5% |
| PFC_EGR1_100K_1 | 9285737 | 91% | 8104052 | 87% | 6212 | 10.7% |
| PFC_EGR1_100K_2 | 8557087 | 51% | 7510446 | 88% | 6921 | 11.0% |
| Cerebellum_EGR1_100K_1 | 9882524 | 95% | 8739064 | 88% | 17795 | 16.2% |
| Cerebellum_EGR1_100K_2 | 13141514 | 95% | 11799729 | 90% | 22866 | 15.4% |

| | | | | | | |
|---|---|---|---|---|---|---|
| PFC_MEF2C_100K_1 | 8478934 | 82% | 7497951 | 88% | 1354 | 7.5% |
| PFC_MEF2C_100K_2 | 11407867 | 91% | 10012928 | 88% | 5362 | 9.1% |
| Cerebellum_MEF2C_100K_1 | 11268711 | 95% | 10016208 | 89% | 24063 | 19.0% |
| Cerebellum_MEF2C_100K_2 | 11481247 | 95% | 10025093 | 87% | 17393 | 15.6% |
| PFC_HDAC2_100K_1 | 14011639 | 85% | 11164030 | 80% | 2929 | 13.6% |
| PFC_HDAC2_100K_2 | 15354891 | 85% | 12429080 | 81% | 3465 | 12.6% |
| PFC_NeuN+_HDAC2_80K_1 | 12258720 | 80% | 10188120 | 83% | 1382 | 11.2% |
| PFC_NeuN+_HDAC2_80K_2 | 12448796 | 80% | 10639071 | 85% | 2842 | 9.6% |

**Supplementary Table S2. List of genes with significant difference in pausing index between PFC and cerebellum based on mouse brain Pol II-total data (fold change > 3, minimal read density > 0.02)**

| PFC < Cerebellum | PFC > Cerebellum |
|---|---|
| Igfbp6 | Dlk2 |
| Plk2 | Rps6ka1 |
| Npas4 | Mycl1 |
| Rec8 | Carhsp1 |
| Snapc2 | Shank3 |
| Kcnf1 | Sparcl1 |
| Nsg1 | Fgfbp3 |
| Slc1a2 | Slc16a8 |
| Grik5 | Gpsm3 |
| Camk2n1 | Smtn |
| Sox12 | Ankrd33b |
| Paqr6 | Fosb |
| Dos | Trip6 |
| Chtf18 | Flrt2 |
| Sept5 | Cul5 |
| Tro | Gm11696 |
| Barhl2 | Gp5 |
| Gm2506 | Cbln1 |
| Syt7 | E130303B06Rik |
| Zfp296 | Htr1b |
| Gpr123 | Fam101a |
| Arpp19 | Proca1 |
| Nfyb | Ccnt1 |
| Rapgefl1 | Pak6 |
| Bex2 | Gpt |
| Lmo4 | Lrg1 |
| Arc | Btbd17 |
| Celf5 | Pnrc1 |
| Tubb2a | Ascl1 |
| Ephx4 | Aqp5 |
| Tmem151a | Kank2 |
| Nap1l1 | Tcte1 |
| Gpc1 | Eif4e |
| Ino80e | 2610203C20Rik |
| Gnmt | Chrm4 |
| Kif17 | Ramp2 |
| Emilin3 | A230052G05Rik |
| Mtch1 | 0610007L01Rik |

| | |
|---|---|
| Arpp21 | Hrc |
| Ptprn | Doc2a |
| Rgs2 | Rsph9 |
| Kcnt1 | Npy1r |
| Hrh3 | Eno3 |
| Stk19 | Scn3b |
| BC018242 | Ntn3 |
| Klhdc8a | Ndrg4 |
| A930018M24Rik | Shroom1 |
| Sun2 | Dusp18 |
| Prkag1 | Loxl3 |
| Pigz | Ttc19 |
| Atp2b1 | Dnaja3 |
| Sstr3 | Def6 |
| Zdhhc8 | Plch2 |
| Man2a2 | Bdh1 |
| Tlcd2 | Nol3 |
| Rgs4 | Coch |
| Ppp1r1a | Gm20537 |
| Micall1 | Acta1 |
| Rasd1 | Tmem88b |
| Eid2 | Cox6b2 |
| Tacc1 | Fam164c |
| Rtn3 | BC106179 |
| Ncan | Fosl1 |
| Bai2 | S1pr5 |
| Gm996 | Ntpcr |
| Gria2 | Msl3l2 |
| Trim9 | Arpc1b |
| Arhgdia | Slc16a3 |
| Nrn1 | Pqlc3 |
| Srm | 1700019N12Rik |
| Vipr1 | Set |
| Gm10116 | Kank3 |
| Fam49a | Mad2l2 |
| Gm4535 | Acyp1 |
| Ttyh3 | |
| BC025920 | |
| Gnao1 | |
| Ltk | |
| Ywhaz | |
| Xrcc3 | |
| Tceb2 | |

| |
|---|
| Sh3glb2 |
| 1700088E04Rik |
| Camk2n2 |
| Arpc2 |
| C030046I01Rik |
| Cacnb1 |
| 1110008P14Rik |
| Gal3st3 |
| Usp30 |
| Ccdc157 |
| Dpysl2 |
| Trim46 |
| Atp1a3 |
| R3hdm1 |
| Eme1 |
| Aifm3 |
| Asphd1 |
| Mcl1 |
| Ppil2 |
| Josd1 |
| Cdc42ep3 |
| Mcrs1 |
| Bloc1s2 |
| Rhog |
| Klhdc8b |
| BC017643 |
| Rnf139 |
| Jdp2 |
| 4931428F04Rik |
| Slc4a3 |
| Rabggta |
| Psenen |
| Efhd2 |
| Pcgf2 |
| Rab35 |
| 1700019L03Rik |
| Hexdc |
| Tcea2 |
| Dusp7 |
| Banf1 |
| Ccdc103 |
| Pdlim7 |
| Pacsin3 |

| |
|---|
| Epn3 |
| Sepw1 |

**Supplementary Table S3. Primers used in qPCR enrichment tests of nMOWChIP-seq libraries.**

| **Species** | **Locus name** | **Primer** |
|---|---|---|
| Human (GM12878 cells) | *Actg1* (F) | CGG AAA GAT CGC CAT ATA TGG AC |
| | *Actg1* (R) | ACC GGC AGA GAA ACG CGA |
| | *Polr2a* (F) | GAG AGA CAA ACT GCC GTA ACC |
| | *Polr2a* (R) | GGG AAA TAA GGA GCG AAA GGA G |
| | *Afm* (F) | GCA GAA CCT AGT TCC TCC TTC AAC |
| | *Afm* (R) | AGT CAT CCC TTC CTA CAG ACT GAG A |
| | *Kctd16* (F) | GCC ATA AAG AAA CAC ACA TGG AAA C |
| | *Kctd16* (R) | GCA AAG CAG GAA CCC TCA AA |
| Mouse brain (PFC and cerebellum) | *Arc* (F) | CAG CAT AAA TAG CCG CTG GT |
| | *Arc* (R) | GTC GCC GCT GAA GCT AGA |
| | *Polr2a* (F) | AAA GAA GGG AGG AGA GGA GGA |
| | *Polr2a* (R) | GGG AGA GAC AAA CTG CCG TAA |
| | *Afm* (F) | CAT TTG ACC CAA ACT GCT AAG T |
| | *Afm* (R) | GTA TGT GAA GTG TCA GCA ATG G |
| | *Gcg* (F) | TCA ACC CAA AGT CCC TGA AG |
| | *Gcg* (R) | CCA AGA GGT TGC ATT GGA AG |
